# Does the evolution of predatory behaviour alter intermale aggression? Insights from a selection experiment on bank voles

**DOI:** 10.64898/2026.06.16.732607

**Authors:** Gokul Bhaskaran, Natalia Boron, Paweł Koteja, Edyta T. Sadowska

**Author notes:** Corresponding author Name: Gokul Bhaskaran, Address: Gronostajowa 7, 30-387 Krakow, Poland. Email addresses: Gokul Bhaskaran, Natalia Boron, Paweł Koteja, Edyta T. Sadowska.

## Abstract

Aggression occurs in many forms and can be an important adaptive behaviour. Two distinct forms are predatory and intermale aggression. It remains unclear whether they share genetic and neurobiological regulatory mechanisms and thus whether evolution of one may influence the other. We tested whether selection for increased predatory behaviour leads to increased intermale aggression using an experimental evolution model comprising lines of bank voles (*Clethrionomys = Myodes glareolus*) selected for high predatory propensity towards crickets (P lines) and unselected control lines (C lines). Adult males were tested in a cricket-hunting test followed by two intermale aggression tests. As expected, P-line males showed higher hunting propensity and performance than C-line males. In the intermale aggression test, a greater proportion of P-line males displayed aggressive behaviours (93% vs. 80%), they did it earlier (mean±SD: 116 ± 144 s vs. 349 ± 320 s), more frequently (33 ± 38 vs. 12 ± 17), and for longer (92 ± 137 s vs. 31 ± 52 s). P-line males also showed a proactive behavioural profile, whereas C-line males were vigilant, spending more time observing the opponent and staying immobile. These results indicate that predatory and intermale aggression partly share genetic and neural regulatory mechanisms.

## 1. Introduction

Aggression comprises a broad spectrum of behaviours, varying in origin, underlying motivations, neurobiological mechanisms, forms of expression, and functional significance (Austerman, 2017; Miczek et al., 2007; Siever, 2008). Among these, predatory and intermale aggression are often regarded as particularly distinct forms (Nelson, 2005; Zhao et al., 2023). Predatory aggression, in which a predator attacks, kills, and often consumes prey, is essential for resource acquisition and survival (Ramirez, 1998; Siegel et al., 1999). In contrast, intermale aggression is directed towards conspecific male competitors and often involves ritualised displays, threats, and physical confrontations that determine social dominance and access to mates and other resources (Anpilov et al., 2025). Notably, there are arguments that the characteristics and neurophysiological mechanisms of predatory behaviour differ so fundamentally from other forms of aggression that it should not be classified as aggression at all (Nelson, 2005; Ramirez, 1998). Nevertheless, understanding whether these distinct forms of aggression are evolutionarily or mechanistically linked provides insight into how complex behavioural traits coevolve under selection. Here, we asked whether selection for enhanced predatory behaviour also increases intermale aggression in lines of a non-laboratory rodent, the bank vole (*Clethrionomys = Myodes glareolus*), selectively bred for increased predatory behaviour towards crickets.

The relationship between predatory and intermale aggression is not straightforward. Barr et al. (1975) found that muricidal rats exhibited more intermale fighting than non-killers, which implies the presence of common underlying mechanisms. However, other studies in rodents have found little or no evidence for such a relationship (Brain & Al-Maliki, 1978; Butler, 1973; Ferrari et al., 1996; Lynds, 1980). The brain circuits supporting these behaviours also appear partly distinct, with rivalry aggression primarily regulated by the medial amygdala and predatory aggression by the central amygdala, while abnormal forms of violence involve overlapping but dysregulated activation of both regions (Haller, 2018). Injection of the same dose of a serotonin agonist into outbred mice significantly reduced intermale aggression, but did not affect predatory aggression (Ferrari et al., 1996). Although regulation of serotonin has been strongly implicated in the control of intermale aggression (Nelson & Chiavegatto, 2001) and other studies suggest a similar control of predatory aggression (Nikulina, 1991; Pucilowski & Kostowski, 1983; Summers et al., 2005; Szebik et al., 2025), it remains unclear exactly how serotonin interacts with the neuronal circuitry underlying different forms of aggression. Taken together, these findings indicate that the motivational and neural dissociations between predatory and intermale aggression underline the complexity of their relationship.

While intermale aggression has been studied widely, the neurophysiological basis of predatory aggression remains less explored. This is notable because, in several respects, predatory aggression resembles the callous–unemotional form of conspecific aggression - a pathological type of aggression characterised by low emotional arousal, lack of empathy, and reduced sensitivity to social cues, as observed in some mammals including humans (Siegel & Victoroff, 2009). These forms of aggression share similar physiological, neurobiological, and behavioural features (Gregg & Siegel, 2001; Haller, 2018; Haller et al., 2001; Tulogdi et al., 2015). There is also evidence suggesting that antisocial and predatory aggression are not only similar in their expression but are regulated by partially overlapping neural mechanisms (Tulogdi et al., 2010, 2015). Hence, studying the mechanisms underlying extreme predatory behaviour in animals can shed light on the neural basis of aberrant conspecific aggression, providing insights into how different forms of aggression are interrelated and regulated across species.

Experimental evolution offers a resourceful tool for uncovering and studying the genetic and regulatory links between the behavioural traits (Rhodes & Kawecki, 2009). A correlated change in the predatory aggression in response to the direct selection for the intermale aggression, or *vice versa*, would imply the presence of a shared genetic background and regulatory mechanisms. However, a systematic literature review (Table S1) revealed that no study has directly tested whether selection for predatory behaviour affects intermale aggression and that only one study has tested the opposite (Sandnabba, 1995a). This single study showed that divergent selection for high or low intermale aggression in laboratory mice resulted in an increased predatory behaviour in males but not in females. A few other selection experiments targeting different traits have provided only indirect insights. Selection for reduced fear-induced aggression in rats did not affect intermale or predatory aggression, suggesting that fear-induced aggression is subject to distinct genetic control (Popova et al., 1993). Laboratory mice from lines selected for high voluntary wheel running, characterised by an altered dopamine signalling, exhibited increased predatory aggression compared to those from non-selected control lines (Gammie et al., 2003). However, testosterone level was not affected by the selection and the high-runner mice tended to show a decreased male dominance in a tube test. These results suggest that the two forms of aggression are controlled by distinct hormonal mechanisms. In addition to ours, we are aware of only one other selection experiment that has directly targeted predatory behaviour. Selection for predatory behaviour towards locusts in golden hamsters (*Mesocricetus auratus*) affected several other traits, but intermale aggression was not studied (Polsky, 1974, 1977, 1978). Our study aims to address this gap by investigating whether artificial selection for enhanced predatory behaviour in bank voles results in correlated changes in intermale aggression.

We used a unique rodent model system comprising lines of bank voles selectively bred for the ability to rapidly capture live crickets in a 10-minute test following a short fasting period (P – Predatory lines), and unselected Control (C) lines (Sadowska et al., 2015). The effects of selection were apparent from the third generation onward with a higher proportion of P-line voles catching crickets, and doing so faster than C-line voles (Fig. S1). Voles from P lines not only show predatory behaviour in the laboratory cricket hunting test, but also prefer animal-based food under semi-natural conditions, as evidenced by their higher hair δ^15^N values than C lines (Hämäläinen et al., 2022), indicating that selection has led to the evolution of a predatory lifestyle. They also show several other characteristics of active hunting predators, including higher food consumption and increased home cage activity (Koteja et al., 2009). Moreover, compared to C-line voles, the P-line ones exhibit proactive personality traits, such as increased activity and boldness, and moving faster on straighter trajectories (Maiti et al., 2019). The evolution of a predatory lifestyle in P-line voles has been associated with increased expression of genes involved in the biosynthesis of testosterone, serotonin and key neurotransmitters (Konczal et al., 2016). As mentioned earlier, testosterone and serotonin have been reported to mediate conspecific aggression in rodents (Nelson & Chiavegatto, 2001), and other studies suggest a similar control of predatory aggression. Together, these characteristics make this selection model well-suited for testing whether evolutionary changes in predatory behaviour lead to correlated shifts in intermale aggression.

We hypothesised that the enhanced predatory tendency in P-line voles is accompanied by increased intermale aggression. To test this hypothesis, we quantified the predatory aggression in cricket-hunting tests (as applied in the selection procedure), and intermale aggression in standardised male-male contest tests (in which two unfamiliar males are placed in a neutral arena). We predicted that P-line males would display higher aggressive propensity, expressed as shorter approach latencies and more frequent and persistent aggressive interactions.

## 2. Materials and methods

### 2.1 Animals and the selection experiment

This work was performed on bank voles (*Clethrionomys = Myodes glareolus* Schreber 1780), a common rodent frequently used as a model in biological research (Górska et al. 2025), from generation 36 and 38 of an ongoing artificial selection experiment (Sadowska et al., 2008, 2015). The rationale, history, and protocols of our selection experiment has been described in details elsewhere (Lipowska et al., 2020; Sadowska et al., 2008, 2015). Briefly, the colony was established with about 320 wild voles caught from Niepołomice Forest in southern Poland during 2000 and 2001. Following five to six generations of random breeding, four replicate lines selected for predatory propensity (Predatory - P) and four randomly bred unselected Control (C) lines were established, with 15-20 reproducing families per line.

Animals were housed in standard opaque polypropylene mouse cages with sawdust bedding, under a constant temperature (20 ± 1°C) and photoperiod (16 h : 8 h light : dark; light phase starting at 3 am). Breeding pairs and pairs with offspring (up to 17 days old) were maintained in model 1290D cages (Tecniplast, Bugugiatte, Italy; dimensions L×W×H: 425×266×155 mm, floor area 800 cm^2^), equipped with a shelter, additional nest material (paper towels) and cardboard tubes for environmental enrichment. Offspring were weaned at 17 days of age, temporarily marked by fur clipping, and kept in family groups until 30–35 days old. At the age of about 34 days, all individuals were permanently marked using mouse ear tags (model 10005-1; National Band and Tag, Newport, KY; mass 0.18 g) and then housed in same-sex trios in model 1264C cages (dimensions L×W×H: 267×207×140 mm; floor area 370 cm^2^). Cages were cleaned about once per week, depending on cleanliness. Water and food (a “breeding type” rodent chow: 24% protein, 3% fat, 4% fibre; Labofeed H, Kcynia, Poland) was provided *ad libitum*.

The predatory propensity was assessed in a cricket hunting test performed on adult animals (75-105 days of age), fasted before the trials. After fasting for a few hours, a live cricket (*Gryllus assimilis*) was introduced to the cage and its presence and state were checked at intervals of 0.5, 1, 3, 6 and 10 min. The results were scored as ranks (rank 1–5: cricket caught in 0.5, 1, 3, 6, or 10 min, respectively; rank 6: cricket not caught). In the early generations the voles were fasted for 10-12 hours before the tests, and the crickets were instars of about 1 cm length. In subsequent generations the fasting duration was gradually decreased (eventually to 2 hours), and the size of the crickets increased to about 2 cm. In the first two generations, three independent tests were performed (each on a different day). Later, the tests were performed on two days (with 7-10 days intervals), but two tests were performed on each day: one during light phase (about 15:00-16:00 hours) and the next after the beginning of dark phase of the colony photoperiod (18:00-19:00 hours), but with lights on. Further details of selection protocol are detailed in Sadowska et al. (2015). Selection was based on the average rank score across trials, adjusted by ANCOVA for sex, body mass, and other cofactors.

### 2.2 Experimental design

This experiment was part of a larger project, whose overall design is shown in Fig. S2. The animals used in this project remained in their natal family groups until ear-tagging at 32 days of age. After tagging, the focal males were housed individually to prevent the formation of dominance hierarchies (Kruczek & Zatorska, 2008).

For this part of the project, we aimed to obtain data from at least 80 males (10 per replicate line) that successfully completed three behavioural tests: the cricket hunting test and two intermale aggression trials. Because some exclusions during the experimental procedures were anticipated, larger initial cohorts were used. During the intermale aggression tests, one C-line male escaped and five males (one C-line and four P-line) were excluded because the tests had to be terminated due to visible bleeding of the opponent males caused by excessive aggression of the focal males. Consequently, final sample included 44 P-line and 45 C-line males (19 C-line and 20 P-line males from generation 36, and 26 C-line and 24 P-line males from generation 38, with 10-12 males per replicate line, 1-2 males per family).

Wherever possible, animals from the first litters were used. However, when the number of available males from first litters was insufficient to achieve the intended representation across replicate lines, additional males from second litters were included. Consequently, the final sample included 40 P-line and 37 C-line males from first litters and only 4 P-line and 8 C-line males from second litters.

At 89–109 days of age (average: 95 days), males were paired with adult females from the same selection line. Eight days post-mating the animals underwent a cricket hunting test, after which they were returned to their home cage. On the 11^th^ and 13^th^ days after mating, focal males were separated from their mates and subjected to the first and second intermale aggression tests, respectively. Following the first trial, the males were returned to their mates and housed under standard colony conditions. After the second trial, the males were euthanised and used in subsequent analyses as part of the larger project (Fig. S2). For logistical reasons, mating and subsequent behavioural testing were staggered across several days, resulting in one to four intermale aggression tests being conducted per day.

### 2.3 Cricket hunting test

The hunting test was conducted during the daytime between 14:00 and 19:00 Central European Summer Time (CEST). The male and its mate were weighed (± 0.01 g), and subjected to a 2-hour fasting period with water available *ad libitum.* After fasting, the male was transferred to a test arena located in the video-recording room. The test arena measured 50 × 50 × 50 cm (length × width × height). To prevent the vole from settling in the corners, each corner was sealed with panels made of the same material as the arena. The male was given a 3-min habituation period, after which a live cricket (imago, approximately 2 cm long) was introduced for a 10-min test. After the test, the focal male was returned to its home cage with the female and provided with food and water. The cage was then transferred back to the animal room.

### 2.4 Intermale aggression tests

The intermale aggression test design followed the general framework of the “standard opponent” paradigm (Brain & Al-Maliki, 1978; Kruczek, 1997; Kruczek & Zatorska, 2008; Marchlewska-Koj et al., 1989; Radwan et al., 2004), commonly used to assess dominance or intermale aggression. However, unlike in some of the previous studies, we used adults rather than juveniles as opponents. This allowed us to standardise opponent characteristics across trials and avoid variation associated with developmental stage, as would be expected when using juveniles that differ even slightly in age. The tests were conducted in a neutral arena to minimise the influence of territoriality (Brain & Al-Maliki, 1978; Kruczek & Zatorska, 2008; Radwan et al., 2004).

The tests were conducted during the daytime, between 08:00 and 14:00 CEST. In each trial, the focal male was tested against a sexually naïve adult opponent male from the C-lines, but originating from a different replicate line than the focal male. Opponent males had no prior exposure to crickets and were housed individually under the same laboratory conditions. To avoid potential familiarity effects (Parmigiani & Brain, 1983), each opponent male was used only once.

On each trial day, the focal male was separated from his female partner and weighed. The opponent male was also weighed, and both males were simultaneously placed into a test arena of the same type as that used in the hunting test for a 20-minute trial. Opponent males were unfamiliar with such arenas. Importantly, both males were introduced into the arena simultaneously, ensuring that neither individual had prior ownership of the test space. Consequently, while the arena was not entirely neutral in terms of general familiarity for focal males, it was neutral with respect to territoriality. This design was intended to evoke general aggressive responses rather than territorial defence, allowing a clearer assessment of behavioural differences shaped by selection.

To distinguish focal from opponent males in the video recordings, a small patch of fur was shaved from either the dorsal neck region or from the rump region of the body. Shaving assignments were balanced across individuals to avoid any systematic effect of shaving location on behaviour. Shaving was performed one day prior to the first intermale aggression trial. For the second trial, additional shaving was not required, as the original marking remained clearly visible.

### 2.5 Video recordings and behavioural analysis

Both the predatory and intermale aggression tests were video-recorded using an analogue camera (SAMSUNG SCB-3000P; resolution: 704 × 576 pixels; 25 frames/sec) mounted 250 cm above the arena floor. The camera was connected to a digital recording system (BCS, Warsaw, Poland) operated remotely from outside the testing room. A monitor screen enabled real-time supervision of the trials.

Each video recording was assigned with a unique ID and analysed using BORIS software (Friard & Gamba, 2016), by an observer blind to the selection line origin of the focal male. Behavioural traits were classified as either point events (for which only frequency and latency were recorded) or state events (for which frequency, latency, and duration were available). Latency was defined as the time from the start of the trial to the first occurrence of a given behaviour, irrespective of the behaviour of the stimulus animal (the cricket or opponent male).

Information about all recorded behaviours is summarized in Table S2. In both the cricket hunting and intermale aggression tests, we quantified approach, resting, grooming, jumping, observing, and exploration. The behaviours were defined as follows: approach, directed movement towards the stimulus animal; resting, periods during which the vole remained immobile and was not engaged in any other observable behaviour; grooming, self-directed maintenance behaviours such as licking, scratching, or cleaning of the fur or body; jumping, sudden vertical or rapid movements directed towards cage walls or boundaries, rather than specifically towards the stimulus animal; observing, a stationary state in which the vole visually attended to the stimulus without approaching or initiating interaction.

Exploration, a general locomotor and exploratory activity, was not directly annotated during video scoring. Instead, exploration duration was calculated as the total test duration (600 s in the cricket hunting test and 1200 s in the intermale aggression test) minus the summed durations of all annotated state behaviours.

In the hunting test, we recorded latencies to spot, approach, and initiate hunting of the cricket, as well as hunting duration, capture success, and time to capture. Spotting, i.e. the initial detection of the cricket, was operationalised as the first clear orientation or visual attention towards the prey. Hunting was defined as an active prey-directed behaviour, including pursuit, rapid orientation, chasing, or lunging towards the cricket. Hunting duration was the total time spent in such behaviour, irrespective of capture success. Capture was defined as successful when the vole grasped and killed the cricket, and time to capture as the time to grasping the cricked followed by killing it.

In the intermale aggression tests, behavioural scoring followed established ethograms used in studies of social interactions and aggression in rodents (Clarke, 1956; Kruczek, 1997; Kruczek & Zatorska, 2008; Marchlewska-Koj et al., 1989; Miczek et al., 2001; Miller et al., 2023; Radwan et al., 2004). We recorded latencies, frequencies, and durations of chasing, wrestling, boxing, escaping, olfactory investigation, and postural displays (supine and squatting) for both the focal and opponent males.

Chasing was defined as rapid directed pursuit of one vole by another; boxing as an upright interaction involving forelimb striking or pushing; wrestling as close-contact physical engagement such as grappling or rolling; and escaping as movement directed away from another individual. These behaviours are well-established components of rodent agonistic repertoires (Clarke, 1956; Miller et al., 2023). We further recorded the latency to the first aggressive behaviour expressed by a given individual, irrespective of which individual initiated the aggressive interaction, and computed duration and frequency of “all aggression” as the sums of durations or frequencies of chasing, boxing, and wrestling.

Nosing (or olfactory investigation) was defined as sniffing or close-contact investigation using the nose directed towards the environment or another individual. Postural behaviours included the supine posture, in which a vole lay on its back for at least three seconds, typically associated with submissive or defensive responses (Atrooz et al., 2021; Patki et al., 2015), and the squatting posture, defined as a low crouched position with reduced movement, often reflecting a non-engaging or defensive state (Clarke, 1956).

To provide an overall characterisation of behavioural tendencies during intermale interactions, composite indices were calculated by combining multiple behavioural components reflecting aggression, defence, and behavioural initiative. Because the intermale aggression test did not produce a single unequivocal outcome such as a clear winner or loser, these indices were intended to summarise the overall behavioural profile and relative dominance tendency of the focal male in relation to the opponent.

None of the traits characterising the behaviour of the focal male indicated whether the male was more aggressive than, or dominant over, the opponent. Therefore, two composite aggression indices were calculated based on the behavioural traits of both the focal and opponent males (Table S2). The frequency- and duration-based aggression indices were calculated using standardised (z-score) differences between the focal and opponent males in the log□□-transformed frequency or duration of behaviours describing the intermale interactions: “aggressive” (chasing, wrestling, and boxing) and “defensive” (escaping, supine posture, and squatting posture). Standardization was applied to ensure equal weight of each component. The log transformation was motivated by the distribution of residuals revealed in the statistical analyses (section 2.6). To allow log transformation of durations and frequencies for behaviours not observed in a particular individual, the values were set to assumed “detection limits”: 0.1 s for durations and 0.5 for frequencies. The rationale for the former was that the shortest durations we recorded lasted 0.15 s. The rationale for the latter was that in some cases it was not clear whether a behaviour had occurred, so 0.5 was a conservative representation of the uncertainty between 0 and 1. The indices were then calculated as the sums of the “aggressive” minus “defensive” components. In addition, a proactivity index was calculated in the same way, based on standardised differences between the focal and opponent males in log-transformed durations of exploration and olfactory investigation (positive contributions to the index value), and resting duration and approach latency (negative contributions).

### 2.6 Statistical analyses

All analyses were performed within the framework of generalized linear mixed models using the GLIMMIX procedure in SAS (version 9.4; SAS Institute, Cary, NC, USA). Binary traits were analysed assuming a binomial distribution with a logit link function and estimated using residual pseudo-likelihood, whereas quantitative traits were analysed assuming a Gaussian distribution using restricted maximum likelihood (REML).

In all models, selection direction was included as the main fixed factor. Results from the intermale aggression tests were analysed with repeated-measures models including trial number (first vs second test) as an additional, within-subject fixed factor. Body mass measured before the predatory and intermale aggression tests was analysed with a similar repeated-measures model, but for three trials and without including body mass as a covariate. Generation was included as a cofactor, and body mass, age, test date, and test time were included as covariates in all models; in models for results from the intermale tests, opponent body mass was included as an additional covariate. All covariates were retained irrespective of significance. Litter number was not included as a cofactor because only a few males originated from the second litter. The initial models also included interactions between selection direction and focal male body mass and, in repeated-measures models, between selection direction and trial number. To simplify interpretation of the main factors, the interactions were removed if not significant.

In all analyses, replicate lines nested within selection direction were included as random effects, together with their interactions with the relevant main fixed factors. In order to properly test the main factors, these effects were retained in the models regardless of significance. However, the estimates of certain random effects could be constrained to zero. As the Satterthwaite approximation of degrees of freedom was applied to the tests of the fixed factors, these models are equivalent to models from which these random effects would be excluded.

Prior to the main analyses, preliminary models were fitted to determine the appropriate residual (co)variance structure and test for heterogeneity of variances among individuals. For the results from cricket hutting test, two candidate models were considered: with equal and unequal variances between selection directions. For the repeated-measures analyses of results from intermale aggression test, six candidate models were considered: (i) equal variances across trial number and selection line; (ii) R-side matrix with unequal (co)variances for trial number only; (iii) G-side matrix with unequal variances between the selection directions at the among-individual level; (iv) R-side matrix with unequal variances for selection line at both within- and among-individual levels; (v) combined R- an d G-side matrices with unequal variances across trial number and selection line (at the among-individual level only); and (vi) R-side matrix with unequal variances for both trial number and selection line at both within- and among-individual levels. For the analyses of body mass (repeated-measures model with three repeated trials), the same six models were fitted, but the equality and inequality also concerned residual covariances across pairs of trials. For binary traits, candidate models were compared based on −2 residual log pseudo-likelihood values and dispersion parameters, with preferred models showing lower values and dispersion close to 1. For quantitative traits, models were compared using the Akaike information criterion (AIC), and when competing models showed similar support (ΔAIC < 2), the model with the simpler covariance structure was preferred, following the principle of parsimony.

Model diagnostics were assessed based on the distribution of residuals and commonly used diagnostic plots (residuals vs. predicted values, histograms, and quantile plots). The distributions of most behavioural traits, except observing duration in the intermale aggression test and exploration duration in the hunting and intermale aggression tests, were right-skewed, and the variables were log_10_ transformed to improve the normality and homoscedasticity of residuals (Tables S8, S9). Data points with an absolute studentized residual values exceeding 3.5 were treated as outliers and excluded from the analyses of the respective trait (but other results obtained on the same individual were not excluded).

The analysis of most of the traits was complicated by the fact that not all individuals expressed all behaviours in each trial (Table S3, S4). Consequently, durations and frequencies were zero, whereas latencies were undefined (right-censored at the trial duration). For traits with many zero values, this resulted in (i) zero-inflated distributions for duration and frequency traits and (ii) right-censored distributions and downwardly biased estimates for latency traits. In addition, for most traits the analyses had to be performed on log-transformed values, which excluded observations with zero values even when such cases occurred in only a small number of individuals and the distribution was not markedly distorted.

To resolve these issues, two approaches were applied depending on the number of zero and censored values. First, for traits with relatively few zero or censored observations (intermale aggression test: 1% for approach behaviour and 6% for observing behaviour), which precluded fitting a logistic regression model to analyse occurrence of the behaviours, the zero values were replaced by the detection limits (0.1 s for duration, 0.5 for frequency counts) and undefined latencies by the upper limit (the full trial duration). Second, for traits with a high proportion of zero or censored values (hunting test: 33% for resting behaviour; intermale aggression test: 23 to 49% for nosing/olfactory investigation, resting behaviour, and aggressive behaviours, including chasing, wrestling, boxing, and their sum), a two-part analytical approach was applied: 1) behaviour occurrence (presence/absence) was analysed as a binary trait using logistic regression model; 2) quantitative analyses (assuming normal distribution) were performed only for records in which the behaviour was observed. To provide a basis for overall inferences, p-values from these complementary analyses were combined using Fisher’s method, where the combined test statistic was calculated as χ² = −2 Σ ln(p), with degrees of freedom equal to twice the number of tests, and the corresponding p-value was obtained from the χ² distribution (Sokal & Rohlf, 2013). The method could be applied because the two parts of the analysis are statistically independent.

Descriptive statistics (mean ± SD) for the presence (occurrence), latency, frequency, and duration of particular behaviours in focal and opponent males are provided in the Supplementary Materials (Tables S3–S7). For latency variables, the descriptive statistics were calculated only for individuals displaying the respective behaviour because latency is undefined when a behaviour does not occur. For frequency and duration variables, all observations were included because zero values constitute biologically meaningful outcomes. In the main text, descriptive statistics are reported together with the results of statistical tests concerning model-based estimates (*F*- and *p*-values). These estimates are presented in the figures as adjusted least-squares means (LSMs) with 95% confidence intervals. For analyses performed on transformed variables, the LSMs are presented as back-transformed estimates. Complete model outputs are provided in Supplementary Materials (Tables S8-S10).

## 3. Results

### 3.1 The animal model

Body mass of focal males ranged from 14.8 to 33.8 g (mean ± SD: 23.8 ± 3.2 g). It did not differ between Predatory (P) and Control (C) lines (*F*_1,6.0_ = 0.01, *p* = 0.92, Tables S10) or between the cricket hunting and intermale aggression tests (F_1,114_ = 0.67, *p* = 0.51). None of the cofactors or covariates (generation, age, test date, test time) had a significant effect (*p* ≥ 0.26).

In regular measurements performed as part of the selection experiment in the last several generations (20 - 31), about 95% of P-line voles, but only 10% of C-line voles, captured a cricket at least once in four trials (Fig. S1a-c). In the hunting tests, conducted on animals from generation 36 and 38 before the experiment presented in this study, cricket capture success was on average lower, but remained substantially higher in P lines (about 67%) than in C lines (4%; Fig. S1d).

In the single trial performed as part of the current experiment on males, all voles spotted the cricket and the mean spotting latency appeared shorter in P lines than in C lines (P: 2.4 ± 4.7 s, C: 14 ± 67 s; Table S3). However, this difference was driven by two extreme observations from C lines (studentized residuals exceeding +3.5). After exclusion of these outliers, the mean latency for C lines decreased markedly (0.98 ± 0.68 s) and was significantly shorter than in P lines (F_1,80_ = 6.1, *p* = 0.016, Table S8). None of the cofactors or covariates had a significant effect on spotting latency (*p* ≥ 0.12).

Despite the shorter spotting latency (except for the two outliers), none of the C-line males attempted to hunt the crickets. In P lines, 33 males (75%) attempted hunting, with a mean latency of 76 ± 126 s, and 30 males (68%) captured and killed the cricket, with a mean latency of 107 ± 149 s.

### 3.2 Latency and frequency of approaches

In the hunting test, all but one P-line male (98%), but only 28 C-line males (62%), approached the cricket (mixed repeated-measures logistic regression: F_1,1_ = 9.4, *p* = 0.20). In the intermale test, all P-line males approached the opponent male in both trials, but one C-line male did not approach the opponent in either trial and eight approached in only one trial.

Among males that approached the cricket, P-line males approached more rapidly than C-line males (P: 29 ± 37 s, C: 203 ± 122 s; Table S3; F_1,4.8_ = 45, *p* = 0.001; Table S8; Fig 1a). These results combined with those concerning proportion of individuals that displayed approach showed a highly significant effect of selection (χ² = 17, *p* = 0.002). Males from P lines also approached the opponent males more quickly (P: 52 ± 92 s, C: 191 ± 233 s; Table S5). Mixed-model analyses, with latencies in trials with no approach set to the upper limit (1200 s in the intermale test) confirmed these differences (F_1,82_ = 50, *p* < 0.0001; Table S9; Fig. 1a).

**Figure 1.**
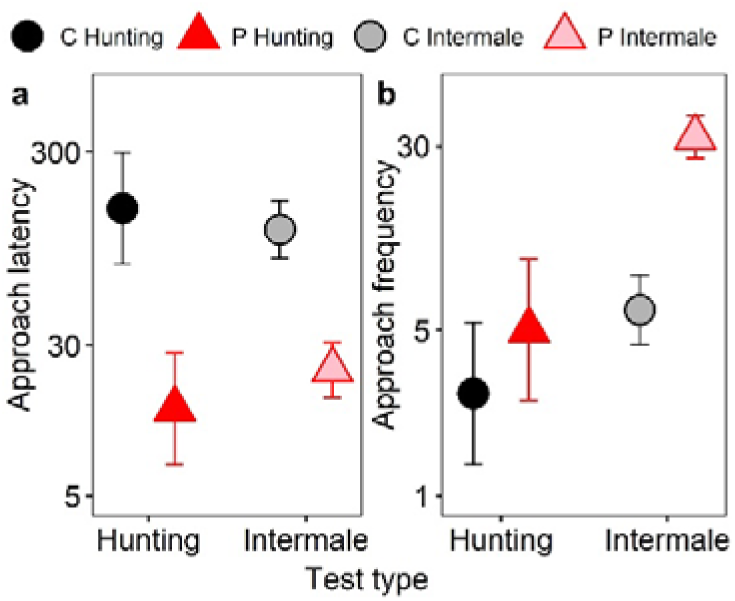
Latency to approach (a; seconds) and frequency of approaches (b; counts) in the cricket-hunting and intermale aggression tests in bank voles from the control (C; n = 45) and predatory-selected (P; n = 44) lines. The values represent adjusted least-squares means (LSMs) with 95% confidence intervals, back-transformed from mixed models fitted to log₁₀-transformed data (y-axes are on a logarithmic scale).

P-line males approached crickets and opponent males more frequently than C-line males (crickets: P: 9.3 ± 9.7, C: 2.1 ± 2.7; Table S3, opponent males: P: 41 ± 34, C: 11 ± 12; Table S6), although the difference appeared statistically significant only for intermale aggression test (hunting test, mixed-model analyses: F_1,6.0_ = 2.5, *p* = 0.17; Table S8; Fig. 1b, and the combined analysis of occurrence and frequency: χ² = 6.8, *p* = 0.15). In the intermale aggression test, with frequencies in trials with no approach set to 0.5 (to allow log transformation), the effect of selection depended on trial number (selection direction × trial number interaction: F_1,60_ = 12, *p* = 0.001), with the difference between the selection lines being greater in the first trial (P: 43 ± 37, C: 8.7 ± 9.5; F_1,107_ = 86, *p* < 0.0001), than in the second trial (P: 40 ± 32, C: 14 ± 13; F_1,107_ = 35, *p* < 0.0001). Nevertheless, P-line males approached the opponent males more frequently overall (F_1,70_ = 75, *p* < 0.0001; Table S9; Fig. 1b). Approach frequency tended to decrease with test time (F_1,99_ = 4.0, *p* = 0.05). None of the other covariates or cofactors significantly affected approach occurrence, latency, or frequency (*p* ≥ 0.054).

### 3.3 Aggressive behaviours in the intermale aggression test

In P lines, 41 out of 44 males (93%) displayed an aggressive behaviour towards the opponent (chasing, wrestling, or boxing) in at least one trial and 34 (77%) did so in both trials, whereas in C lines only 36 of 45 males (80%) displayed aggression in at least one trial and 22 (49%) in both trials (Fig. 2a). The difference was highly significant (mixed repeated-measures logistic regression: F_1,72_ = 7, *p* = 0.009). Moreover, among individuals that displayed the aggressive behaviours, P-line males initiated the aggression more rapidly (latency, P: 116 ± 144 s, C: 349 ± 320 s; F_1,8_ = 20, *p* = 0.002; Table S5; Figure 2e; combined significance from the two parts of the analysis: χ² = 22, *p* = 0.0002). None of the covariates or cofactors affected these characteristics significantly (*p* ≥ 0.12), except that the proportion of aggressive males decreased with test date (F_1,77_ = 4.08, *p* = 0.047).

**Figure 2:**
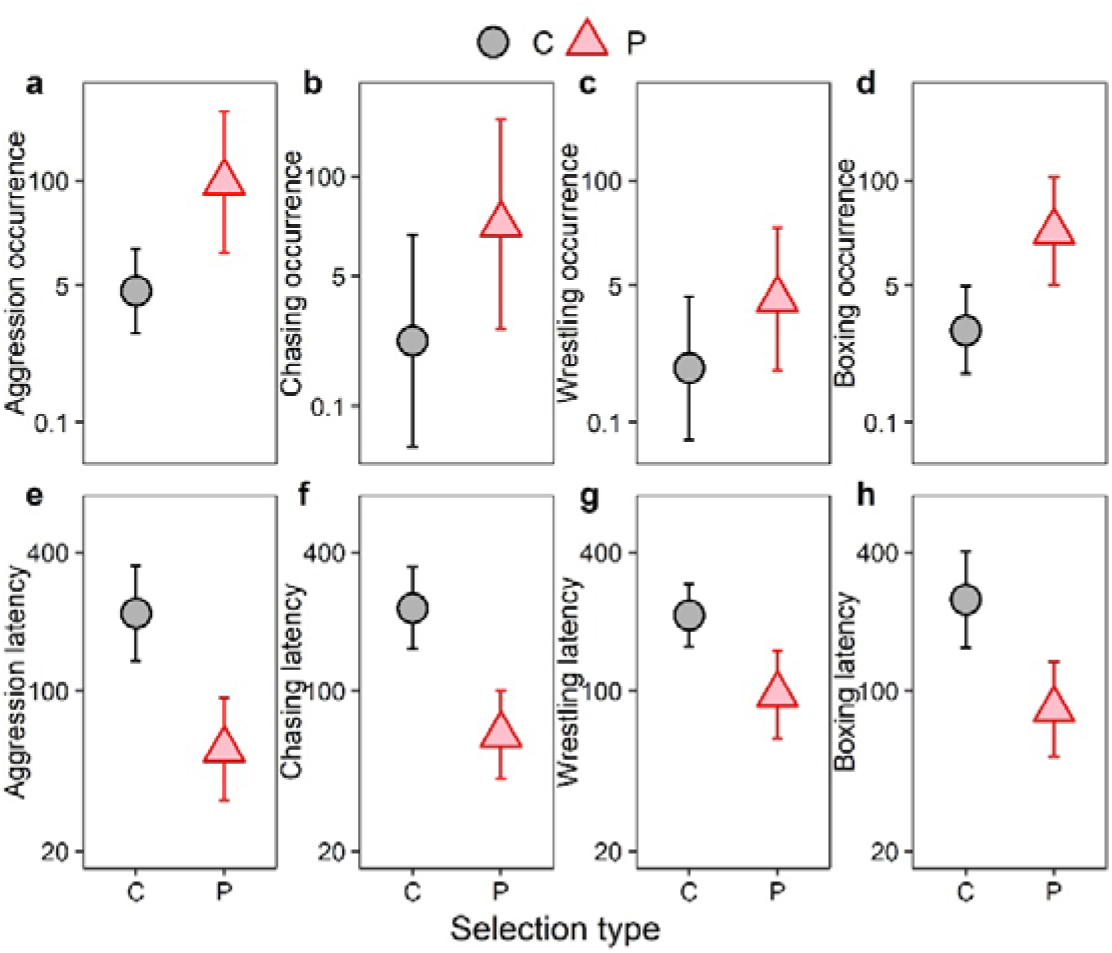
Percentages of males displaying aggressive behaviours (a–d) and latency to the first aggressive behaviour (e–h; seconds) in the intermale aggression test in bank voles from the control (C) and predatory-selected (P) lines. Results are shown for all aggressive behaviours combined (a,e) and for the specific types of aggression: chasing (b,f), wrestling (c,g), and boxing (d,h). The values represent adjusted least-squares means (LSMs) with 95% confidence intervals, back-transformed from mixed models fitted to log₁₀-transformed data (y-axes are on a logarithmic scale).

The results were similar when each of the three types of aggressive behaviour were analysed separately. In P lines, 86% chased the opponent in at least one trial, and 64% in both trials, whereas in C lines 67% in at least one trial and 29% in both trials (F_1,4.7_ = 4.3, *p* = 0.10; Fig. 2b). Similarly, 70% P-line males engaged in wrestling in at least one trial and 50% in both trials, whereas 56% C-line males did so in at least one trial and 29% in both trials (F_1,4.7_ = 3.2, *p* = 0.14; Fig. 2c). Boxing was displayed by 89% P-line males in at least one trial and by 66% in both trials, whereas by 71% C-line males in at least one trial and 38% in both trials (F_1,82_ = 8.2, *p* = 0.005; Fig. 2d). Note, that the magnitude of the effect of selection was similar for each of these traits, and the higher significance of the effect of selection in the case of boxing was mainly due to a smaller random variation among the replicate lines (Table S9).

Among individuals that engaged in the aggressive behaviours, males from P lines engaged in each of the three forms of aggressive behaviours quicker than those from C lines (latency to chasing, P: 133 ± 172 s, C: 322 ± 262 s, F_1,7.4_ = 25, *p* = 0.001; wrestling, P: 209 ± 294 s, C: 300 ± 244 s, F_1,60_ = 8.5, *p* = 0.005; boxing, P: 165 ± 191 s, C: 381 ± 347 s, F_1,6.0_ = 16, *p* = 0.007; Table S5, S9; Fig. 2f,g,h). These results combined with those concerning the proportion of individuals displaying aggressive behaviours showed highly significant effect of selection for each of the three modes of aggression (chasing: χ² = 18, *p* = 0.001; wrestling: χ² = 15, *p* = 0.006; boxing: χ² = 20, *p* = 0.0004). None of the covariates or cofactors significantly affected these traits (p ≥ 0.052), except that the proportion of males displaying chasing behaviour decreased with test time (F_1,162_ = 4.2, *p* = 0.04).

Voles from P lines engaged in the aggressive behaviours more frequently and for a longer time than those from C lines (Tables S6, S7; Fig. 3). The summed frequency of chasing, wrestling, and boxing was 33 ± 38 in P lines and 12 ± 17 in C lines (F_1,5.1_ = 5.4, *p* = 0.06; Fig 3a), and the summed durations were, respectively, 92 ± 137 s and 31 ± 52 s (F_1,5.0_ = 7.8, *p* = 0.03; Fig 3e). These results combined with those concerning proportion of individuals that displayed an aggressive behaviour showed a highly significant effect of selection for both frequency (χ² = 15, *p* = 0.005) and duration (χ² = 16, *p* = 0.003). The aggression frequency was higher in generation 36 than in generation 38 (F_1,67_ = 9.5, *p* = 0.003); no other covariates or covariates had a significant effect on either the frequency or the duration (*p* ≥ 0.08).

**Figure 3:**
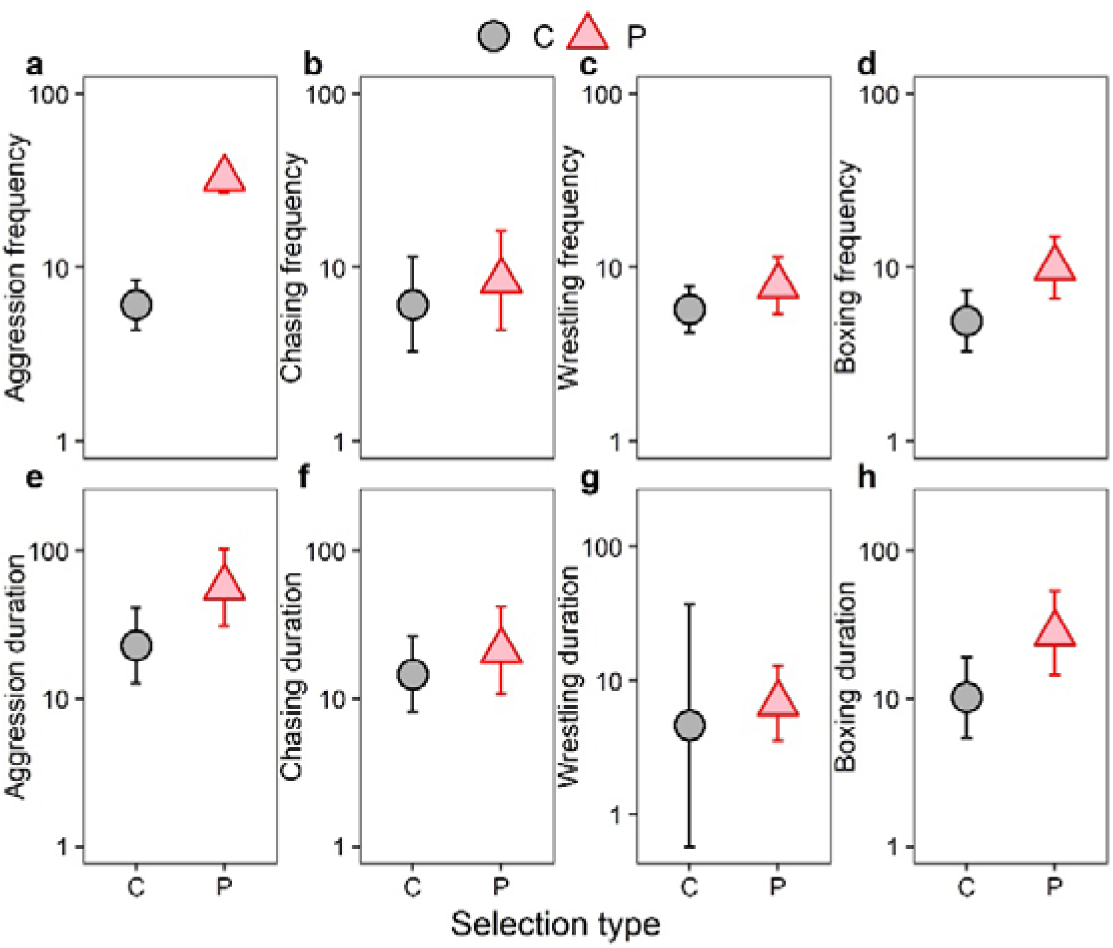
Frequencies (a–d; counts) and durations (e–h; seconds) of aggressive behaviours in the intermale aggression test in bank voles from the control (C) and predatory-selected (P) lines. Results are shown for all aggressive behaviours combined (a,e) and for the specific types of aggression: chasing (b,f), wrestling (c,g), and boxing (d,h). The values represent adjusted least-squares means (LSMs) with 95% confidence intervals, back-transformed from mixed models fitted to log₁₀-transformed data (y-axes are on a logarithmic scale).

Each of the three aggressive behaviours contributed similarly to the magnitude of the difference between P and C lines in the summed frequency and duration of aggressive behaviours. Overall, in animals displaying these behaviours, the frequencies of chasing, wrestling and boxing were similar to each other and were consistently higher in P lines than in C lines (from 2.6 to 2.9 times), although the difference appeared statistically significant only for boxing (chasing: P: 11 ± 14, C: 4.3 ± 6.9, F_1,5.2_ = 0.84, *p* = 0.40; Fig. 3b and wrestling: P: 9.1 ± 13.5, C: 3.5 ± 6.3, F_1,43_ = 1.7, *p* = 0.20; Fig. 3c and boxing: P: 12 ± 15, C: 4.2 ± 6.6; F_1,5.7_ = 9.6, *p* = 0.02; Fig. 3d). For durations the overall pattern was different, because both in P and C lines the duration of wresting was markedly lower, whereas of boxing markedly higher, than the duration of chasing. However, the differences between the P and C lines were similar, and again significant only for boxing (chasing, P: 29 ± 41 s, C: 11 ± 19 s, F_1,3.6_ = 1.6, p = 0.28; Fig. 3f; wrestling, P: 10 ± 17 s, C: 2.9 ± 4.9 s, F_1,2.1_= 1.8, *p* = 0.31; Fig. 3g; boxing, P: 53 ± 121 s, C: 17 ± 42 s, F_1,5.2_ = 8.0, *p* = 0.03; Fig. 3h). These results, combined with those concerning the proportions of individuals engaging in the aggressive behaviours, showed highly significant effect of each of these traits (chasing, frequency: χ² = 6.5, p = 0.17, duration: χ² = 7.2, p = 0.13; wrestling, frequency: χ² = 15, p = 0.006, duration: χ² = 6.3, p = 0.18; boxing, frequency: χ² = 18, *p* = 0.001, duration: χ² = 17, *p* = 0.002).

Chasing frequency decreased with the time at which the test was performed (F_1,31_ = 4.4, *p* = 0.04, Table S9) whereas both its frequency and duration were higher in generation 36 than 38 (F_1,54_ = 16, *p* = 0.0002; F_1,53_ = 5.8, *p* = 0.02, respectively). Wrestling frequency was also higher in generation 36 than 38 (F_1,41_ = 4.9, *p* = 0.03), and it increased with the opponent body mass (F_1,72_ = 15, *p* = 0.0002). None of other covariates or cofactors significantly affected frequencies of or durations of the three aggressive behaviours (*p* ≥ 0.08).

### 3.4 Non-aggressive behaviours in hunting and intermale aggression tests

In the hunting test, observing behaviour occurred in 27 of 44 P-line males (61%) and 44 of 45 C-line males (98%). The difference was significant (mixed repeated-measures logistic regression: cricket: F_1,14_ = 7.4, *p* = 0.02). Among males displaying observing behaviour, P-line males spent less time observing than C-line males (P: 24 ± 50 s, C: 214 ± 157 s; Table S3; F_1,37_ = 57, *p* < 0.0001; Table S8; Fig. 4a). Together, these results indicated a highly significant effect of selection (χ² = 26, *p* = 2.3 × 10^-5^). Observing duration increased with date (F_1,61_ = 5.0, *p* = 0.03) and test time (F_1,64_ = 12, *p* = 0.001). None of the other covariates or cofactors were significant (*p* ≥ 0.15).

**Figure 4:**
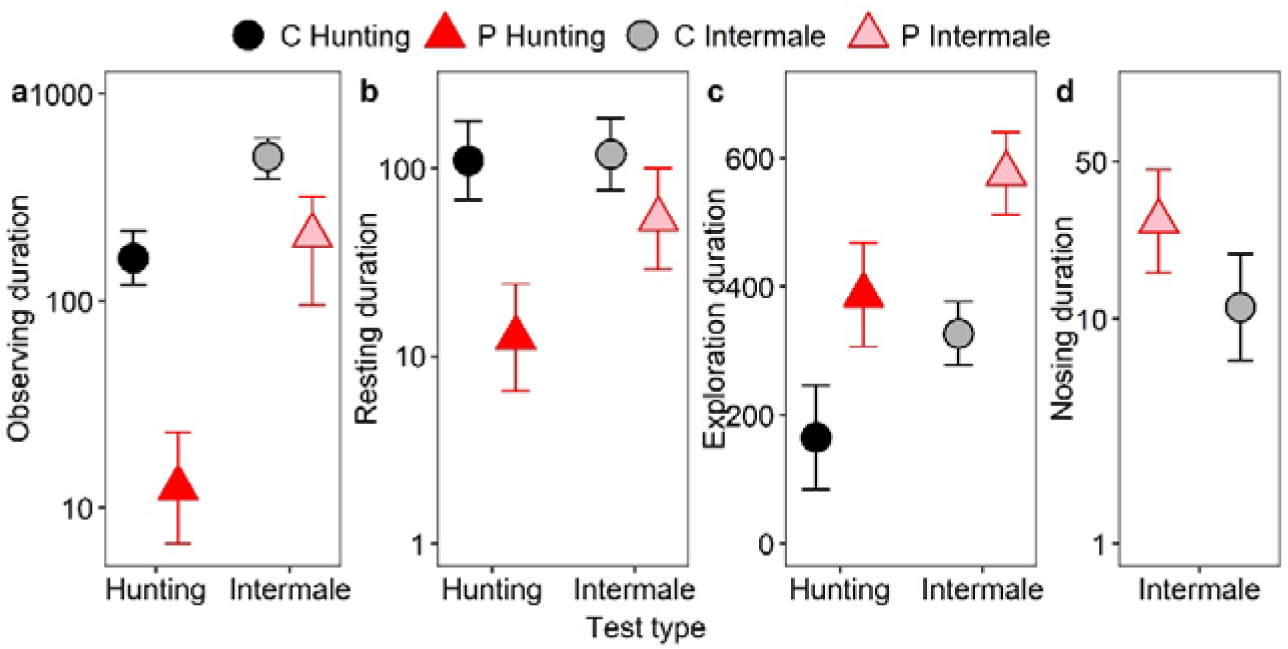
Durations (seconds) of observing (a), resting (b), and exploration (c) in the hunting and intermale aggression tests, and of nosing/olfactory investigation in the intermale aggression test (d), in bank voles from the control (C) and predatory-selected (P) lines. The values represent adjusted least-squares means (LSMs) with 95% confidence intervals derived from mixed models. For observing, resting, and nosing/olfactory investigation, the values are back-transformed from models fitted to log₁₀-transformed data (y-axes on a logarithmic scale).

In the intermale aggression test, all males displayed observing behaviour in at least one trial, whereas 40 P-line males (91%) and 44 C-line males (98%) displayed it in both trials, with P lines spending less time observing opponent males (one observation from P-line with a studentized residual exceeding +3.5 was excluded as an outlier; P: 221 ± 217 s, C: 508 ± 279 s; Table S7). Mixed-model analyses, with zero values replaced by the detection limit (0.1 s), confirmed that these differences were significant (F_1,5.1_ = 22, *p* < 0.005; Table S9; Fig. 4a). Observing duration increased with focal and opponent body mass (focal: F_1,62_ = 5, *p* = 0.02; opponent: F_1,82_ = 5, *p* = 0.03). None of the other covariates or cofactors were significant (*p* ≥ 0.06).

In the hunting test, resting behaviour was displayed by 48% of P-line males and 84% of C-line males. In the intermale aggression test, 61% of P-line males and 82% of C-line males displayed resting behaviour in at least one trial, whereas 30% and 49%, respectively, did so in both trials. The differences were highly significant (mixed repeated-measures logistic regression: hunting: F_1,82_ = 12, *p* = 0.0009; intermale: F_1,99_ = 9.4, *p* = 0.003). In the intermale aggression test, resting behaviour was observed more often in the first than in the second trial (F_1,164_ = 6.3, *p* = 0.01), and in generation 36 than in generation 38 (F_1,102_ = 19, *p* < 0.0001). It decreased with test date (F_1,92_ = 5, *p* = 0.03). None of the covariates or cofactors significantly affected the occurrence of resting behaviour (*p* ≥ 0.06).

Among males displaying resting behaviour, P-line males spent less time resting than C-line males in both tests (hunting: P: 15 ± 43 s, C: 184 ± 183 s; F_1,52_ = 27, *p* < 0.0001; intermale: P: 72 ± 140 s, C: 198 ± 277 s; F_1,53_ = 5.4, *p* = 0.02; Fig. 4b). These results combined with those concerning the proportion of individuals displaying resting behaviours showed highly significant effect of selection (cricket: χ² = 32, *p* = 1.55 × 10□□; opponent: χ² = 19, *p* = 0.0007). None of the other covariates or cofactors significantly affected resting duration (p ≥ 0.09), except that in the intermale aggression test resting duration decreased with increasing opponent body mass (F_1,67_ = 7.5, *p* = 0.008).

Exploration duration was longer in P-line than in C-line males in both tests (hunting: P: 386 ± 133 s, C: 167 ± 151 s; F_1,4.3_ = 27, *p* = 0.006; intermale: P: 574 ± 240 s, C: 326 ± 185 s; F_1,75_ = 39, *p* < 0.001; Fig. 4c). None of the covariates or cofactors significantly affected exploration duration (*p* ≥ 0.11), except that in the intermale aggression test exploration duration decreased with test time (F_1,117_ = 5.8, *p* = 0.02).

Nosing/olfactory investigation was measured only in the intermale aggression test. P-line males spent more time in olfactory activities than C-line males P: 95 ± 204 s, C: 73 ± 183 s; F_1,65_ = 5.3, *p* = 0.02; Fig. 4d). The duration was higher in generation 38 than in generation 36 (F_1,70_ = 4.8, *p* = 0.03), whereas none of the other covariates or cofactors had significant effects (*p* ≥ 0.3).

### 3.5 Indices of intermale aggression and proactivity

The frequency- and duration-based aggression indices, which measure the intensity of aggressive behaviours of the focal male relative to that of the opponent male, were analysed after excluding outliers (frequency: two P-line observations from the first trial with studentized residuals exceeding +3.5; duration: one observation each from the P and C lines in the first trial, with studentized residuals above +3.5 and below −3.5, respectively). Both indices were higher in P-line than in C-line males (Fig. 5a, b), although the difference was not statistically significant for the frequency-based index (mean ± SD: P: 0.55 ± 3.69, C: −0.54 ± 2.31; F_1,70_ = 2.8, *p* = 0.10; Table S9). For the duration-based index, the overall effect of selection was significant (P: 0.61 ± 3.15, C = −0.59 ± 2.65; F_1,166_ = 5.0, *p* = 0.03; Table S9), but it depended strongly on the trial number (interaction: F_1,69_ = 8.4, *p* = 0.005). In the first trial, the index was markedly higher in P-line than in C-line males (P: 1.12 ± 3.40, C: −1.26 ± 2.71; F_1,166_ = 12, *p* = 0.0006), whereas in the second trial the values did not differ between the lines (P: 0.09 ± 2.82, C: 0.08 ± 2.43; F_1,166_ = 0.01, *p* = 0.92; Table S9). The frequency index was higher in generation 36 than in generation 38 (F_1,82_= 4.3, *p* = 0.04) and increased with opponent body mass (F_1,146_ = 4.5, *p* = 0.04). None of the other covariates or cofactors significantly affected either index (*p* ≥ 0.10).

**Figure 5:**
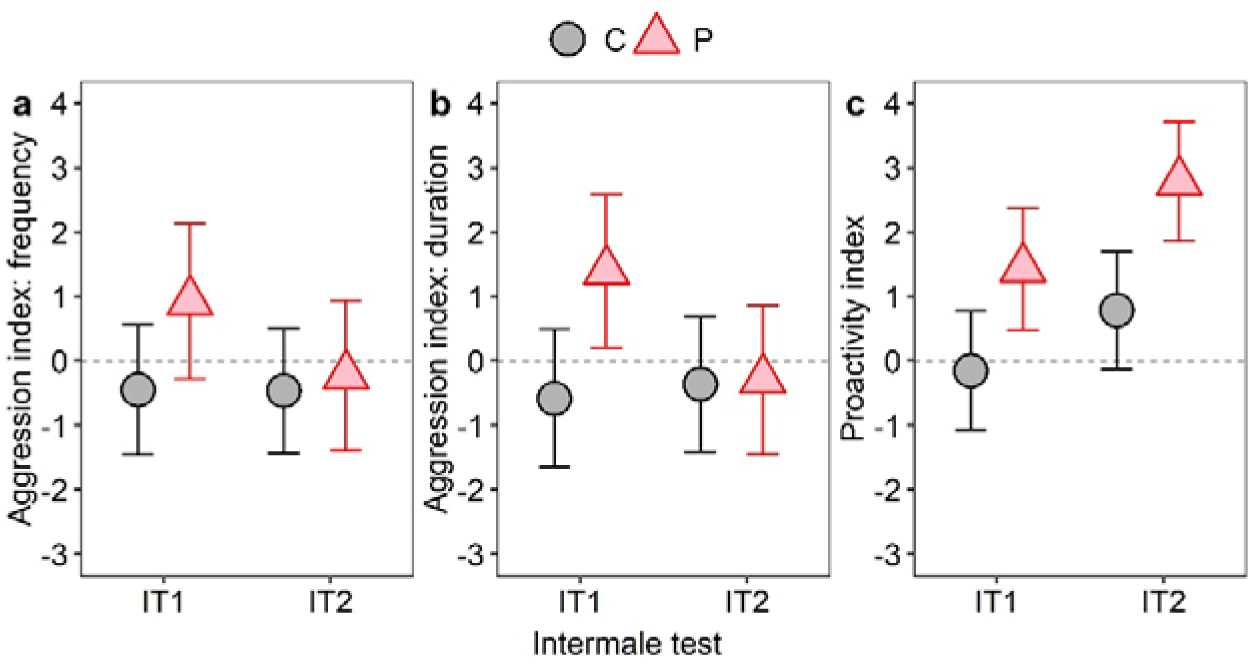
Aggression indexes based on behaviour frequencies (a) and durations (b), and proactivity index (c) in the first (IT1) and second (IT2) intermale aggression tests in bank voles from the control (C) and predatory-selected (P) lines. The values represent adjusted least-squares means (LSMs) with 95% confidence intervals derived from mixed models.

The proactivity index was analysed after excluding one high outlier from a C-line in the second trial. P-line males had a higher proactivity index values than C-line males (P: 2.09 ± 2.12, C: 0.21 ± 2.58; F_1,82_ = 24, *p* < 0.0001; Table S9; Fig. 5c). The index increased with test date (F_1,82_ = 10, *p* = 0.002), whereas none of the other covariates or cofactors significantly affected the index (*p* ≥ 0.12).

## 4. Discussion

Our study demonstrates that long-term selection for predatory behaviour in bank voles resulted in a correlated increase in intermale aggression. Compared with males from the unselected control (C) lines, predatory-selected (P-line) males approached both crickets and opponent males more rapidly and more frequently, were more likely to display aggressive behaviours, initiated aggression sooner, and engaged in aggressive interactions for longer periods. P-line males also spent less time observing and resting, explored more, and had higher proactivity index values, indicating that selection affected a broader behavioural syndrome rather than predatory behaviour alone.

The most direct interpretation of our findings is that predatory and intermale aggression are not entirely independent behavioural traits. Previous studies on the relationship between these forms of aggression in rodents have produced mixed results, with some reporting positive associations (Barr et al., 1975; Sandnabba, 1995a, 1996), and others finding little or no relationship (Brain & Al-Maliki, 1978; Ferrari et al., 1996; Lynds, 1980). Our results support the former view and are consistent with studies of mice selectively bred for high and low levels of intermale aggression, in which highly aggressive lines also showed elevated predatory tendencies (Sandnabba, 1995a, 1995b, 1996). Based on those findings, (Sandnabba, 1996) proposed that the genetic mechanisms underlying predatory and intermale aggression are not entirely distinct. Our results from bank voles provide independent support for that hypothesis.

The correlated response to selection was evident across all forms of intermale aggression. P-line males were more likely to chase, wrestle, and box, and initiated each of these behaviours more rapidly than C-line males. These results suggest that selection for predatory behaviour increased a general propensity to engage in aggressive interactions rather than modifying a specific component of the aggressive repertoire. Although the statistical support was strongest for boxing, the magnitude of the difference between the selection lines was similar for all three behaviours. Thus, the clearer effect observed for boxing likely reflects lower variation among replicate lines rather than a disproportionately strong response of this particular behaviour to selection. Nevertheless, boxing may deserve particular attention because it is often interpreted as a ritualised form of aggression that allows opponents to assess one another and establish dominance while limiting the risk of injury (Clarke, 1956; Grant & Mackintosh, 1963). This upright striking posture may offer both a visual signal and a physical advantage, allowing individuals to assert control while minimising the risk of injury. Supporting this interpretation, Viken & Knutson (1982) found that rats trained with negative reinforcement for aggression displayed significantly more boxing in later resident-intruder encounters, and that these behaviours were often reciprocated by opponents. More generally, these patterns are consistent with the idea that aggressive displays can evolve not only as direct means of conflict resolution but also as signals of competitive ability (Huntingford & Turner, 1987).

The available evidence concerning the role of body mass in establishing dominance relationships is contradictory and appears to vary among rodent species (Hiadlovská et al., 2015; Kim et al., 2015; Nagy et al., 2024). In our study, focal males from the two selection directions did not differ in body mass, indicating that the differences in aggressive behaviour observed here were not a consequence of body size. This finding is consistent with previous work on bank voles, in which dominance was unrelated to body mass (Radwan et al., 2004). The aggression indices provide additional insight into the nature of the behavioural divergence between the lines because they compare the aggressive behaviour of focal males relative to that of their opponents. The aggression indices of C-line focal males were close to zero and, in the case of the duration-based index, even negative. Thus, C-line focal males were, on average, no more aggressive, and sometimes less aggressive, than the opponent males with which they interacted. This observation is important because it suggests that the main results and conclusions do not depend on our choice to using naïve C-line males as opponents.

In contrast, the indices showed that P-line focal males were more aggressive than their opponents. However, the difference between P- and C-line males tended to be greater during the first trial than during the second. This pattern was evident for the aggression indices and, to a lesser extent, for several specific traits.

Previous studies examining the effects of repeated aggressive encounters have produced mixed results. Repeated resident–intruder confrontations in rodents resulted in gradual declines in attack frequency and threat displays, a pattern interpreted primarily as behavioural habituation, although fatigue may also contribute (Koolhaas et al., 2013; Winslow & Miczek, 1984). In contrast, other studies have shown that repeated winning experiences can maintain or even enhance aggression, increasing the probability of success in subsequent encounters through mechanisms involving aggression reward, elevated testosterone, and reinforcement of aggressive motivation (Couppis & Kennedy, 2008; Oyegbile & Marler, 2005; Takahashi, 2022). Our results appear more consistent with the former pattern, although it cannot be explained by fatigue. One possibility is that focal males initially perceived the opponent as a serious threat, but after experiencing that the encounter was temporary, with the opponent subsequently removed and the focal male returned to its home cage, they responded less aggressively during the second trial. Another possibility is that, because the first trial was the first encounter with an unfamiliar male, the P-line focal males escalated the aggression even if the opponent showed signs of submission. Following the experienced gained in the first trial, the focal males may have ceased further aggression once the opponent responded submissively to the initial aggressive interactions. In both cases, the effect of the experience would be less apparent in C-line focal males because, on average, they were no more aggressive than their opponents.

The hunting test revealed an interesting pattern. P- and C-line males did not differ in latency to spot the cricket or in approach frequency. Nevertheless, only P-line males attempted to hunt and successfully captured crickets. Thus, the higher hunting success of P-line males cannot be explained by greater alertness or more frequent encounters with prey. Rather, P-line males were more effective at converting prey encounters into hunting attempts and successful captures.

However, the enhanced aggression of P-line males appears to form part of a broader behavioural syndrome shaped by selection for predatory behaviour. Across both the hunting and intermale aggression tests, P-line males approached stimulus animals more rapidly, spent less time observing and resting, explored more extensively, and had higher proactivity index values than C-line males. Previous work on these lines showed that P-line voles are more active, bolder, and faster to interact with novel objects and environments (Bhaskaran et al., 2026; Maiti et al., 2019), indicating that these behavioural differences are not restricted to the contexts examined here. Similar associations between activity, exploration, and aggression have been reported in other rodents (Hood & Quigley, 2008; Koolhaas et al., 1999), and laboratory mice selected for high voluntary wheel running also exhibit elevated predatory aggression (Gammie et al., 2003). Together, these findings suggest that the correlated increase in intermale aggression observed in P-line voles may arise from broader changes in motivational and behavioural regulation associated with a proactive coping style.

The differences in observing, resting, and olfactory behaviours provide further evidence that P-line males differed from C-line males not only in aggression, but also in their overall behavioural strategy. In both hunting and intermale test, P-line males spent less time observing stimuli and resting, but more time actively interacting with their environment. At first glance, reduced observation may appear paradoxical because successful predation and competitive interactions often require information gathering (Dall et al., 2005; Hämäläinen et al., 2022). However, P-line males also showed higher exploration levels, shorter approach latencies, and, during intermale encounters, more extensive olfactory investigation. Olfactory investigation is a key component of social assessment in rodents, providing information about the identity, physiological condition, reproductive status, and dominance characteristics of conspecifics and influencing the course of aggressive interactions (Galef, 2013). Thus, P-line males did not appear to gather less information; rather, they allocated less time to passive assessment and more time to active engagement compared to C-line males. Such differences are consistent with proactive and reactive behavioural profiles (Koolhaas et al., 1999) and support the idea of correlated evolution of behavioural traits (Réale et al., 2007).

The correlated response observed in our study may reflect shared neurobiological mechanisms underlying predatory and intermale aggression. Studies in rodents and other vertebrates indicate that aggression, social dominance, and behavioural activation are influenced by overlapping neural and neurochemical systems, particularly those involving monoamines (Caramaschi et al., 2008; Chichinadze et al., 2014; Miczek et al., 2007). However, the neurobiological basis of dominance and aggression remains incompletely understood, and it is still unclear to what extent behavioural differences are associated with structural or functional differences in the brain (Kozorovitskiy & Gould, 2004; Wommack & Delville, 2007). Future studies comparing neural, and endocrine profiles of P- and C-line voles could therefore provide valuable insight into the mechanisms underlying the correlated evolution of these behaviours. Although the functions of predatory and intermale aggression differ – prey are subdued and consumed, whereas conflicts between conspecifics usually end in retreat or submission – our results suggest that these behaviours are not entirely independent.

## 5. Conclusions

Our findings demonstrate that long-term selection for predatory behaviour in bank voles resulted in a correlated increase in intermale aggression and a broader proactive behavioural phenotype. These results indicate that predatory and intermale aggression are partially integrated components of behavioural variation and may evolve together through shared underlying mechanisms. Future work examining the neurobiological and endocrine bases of these responses, as well as aggression expressed by females, will help clarify the mechanisms responsible for this correlated evolution.

## Ethics

The animal colony was under the supervision of a qualified veterinary surgeon. All the experimental procedures were approved by the decisions of the Local Bioethical Committee in Cracow, Poland (selection experiment: No. 258/2017 and No. 213/2023; hunting and aggression tests: No. 70/2023).

## Data accessibility

The Supplementary Material document contains Figures S1–S2 and Tables S1–S2, whereas Supplementary Tables S3–S10, together with the raw dataset and associated SAS and R code, are available in an open repository: https://doi.org/10.57903/UJ/A13EOF (Sadowska et al., 2026).

## Declaration of AI use

We have not used AI-assisted technologies in creating this article. Limited assistance was used for language editing, and R codes.

## Authors’ contributions

G.B: Data curation, formal analysis, investigation, methodology, validation, visualization, writing-original draft, writing-review & editing; N.B: Investigation, video analysis; P.K: Conceptualization, formal analysis, funding acquisition, methodology, resources, software, supervision, validation, writing-review & editing; E.T.S: Conceptualization, formal analysis, funding acquisition, methodology, project administration, resources, software, supervision, validation, writing-review & editing.

All authors gave final approval for publication and agreed to be held accountable for the work performed therein.

## Acknowledgments

The authors would like to thank Katarzyna Baliga-Klimczyk, who managed the animal colony. We are grateful to several students for their help with animal maintenance and Konrad Król for his help with the video analyses. We also want to thank Małgorzata M. Lipowska and Alaa Hseiky for their suggestions and insights at various stages of this work.

