## Supplementary materials for "Does the evolution of predatory behaviour alter intermale aggression? Insights from a selection experiment on bank voles"

*Corresponding author

**Corresponding author details:**

Name: Gokul Bhaskaran

**Email addresses:**

**ORCID numbers:**

Gokul Bhaskaran: 0009-0002-1255-4134

Natalia Boron: 0009-0003-9793-9478

Paweł Koteja: 0000-0003-0077-4957

Edyta T. Sadowska: 0000-0003-1240-4814

### Supplementary figures

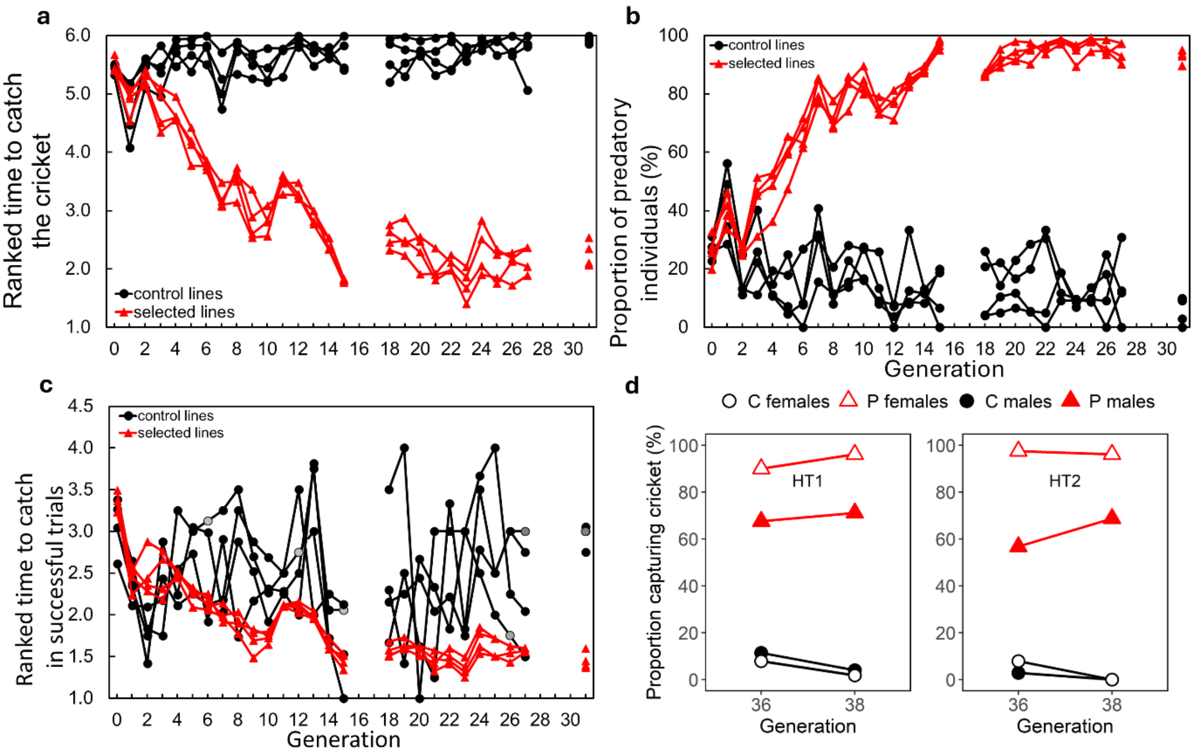


**Figure S1: Panels a-c:** direct phenotypic responses to 31 generations of selection for predatory propensity in bank voles, measured as (a) ranked time to catch a cricket (the selection criterion), (b) proportion of predatory individuals (%), and (c) ranked time to catch a cricket in successful trials (grey points for C lines indicate cases where no individual caught the cricket; these values are estimates based on the mean of the values from the previous and next generation, or on the value from the previous generation when subsequent data were unavailable). The gaps between generations represent periods of relaxed selection. The effects of selection were visible from the third generation onwards. Note that not only the propensity to attack the crickets increased in P lines (b), but also the hunting skilfulness (shorter mean time to catch, computed only for the successful trials; c). **Panel d:** hunting success in the first (HT1) and second (HT2) hunting tests performed on animals from generations 36 and 38 before the experiment presented in this report (see Fig. S2). HT1 was conducted in the animals' standard maintenance cages. HT2 was conducted in a larger test arena for males and in a mating cage for females. Values represent the percentage of individuals that captured and killed a cricket during a single 10-min trial. Data include all animals tested in HT1 and HT2, irrespective of whether they were later included in the present experiment (generation 36: males, C = 35 and P = 37; females, C = 38 and P = 40; generation 38: males, HT1 C = 50 and P = 52, HT2 C = 47 and P = 48; females, C = 54 and P = 52). The proportion of males that captured a cricket was consistently much higher in P than in C lines - (proportions averaged across two generations and two trials, C lines: 4%, P lines: 81%). The proportions observed in the present study were generally lower than those shown in recent generations of the selection experiment (panel b), particularly in the C lines and among P-line males, whereas the proportion of predatory P-line females remained relatively high. This difference may partly reflect the fact that the values shown in panel b represent the proportion of individuals that captured a cricket at least once across four hunting tests, whereas the present estimates are based on performance in a specific hunting test.

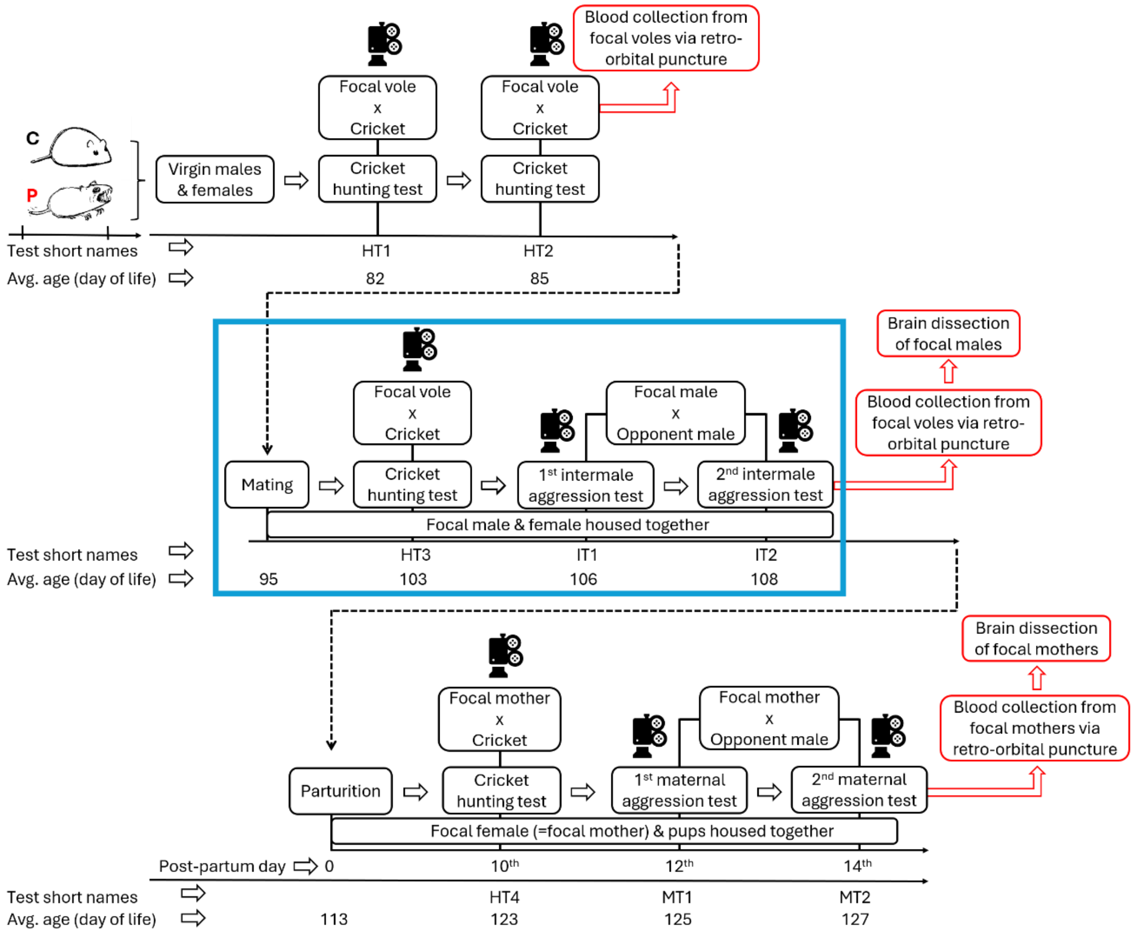


**Figure S2**: Schematic overview of the experimental design conducted as part of a larger project, including predatory (hunting) aggression tests (HT), intermale aggression tests (IT), and maternal aggression tests (MT). The present study focuses on the section highlighted in blue. Before the start of the experiment, virgin males and females from the control (C) and predatory-selected (P) lines were subjected to two hunting tests (HT1 and HT2) (generation 36: 40 P-line males, 40 P-line females, 40 C-line males, and 40 C-line females; generation 38: 50 P-line males, 50 P-line females, 49 C-line males, and 49 C-line females). HT1 was conducted in the animals' standard maintenance cages (Tecniplast 1264C model). HT2 was conducted in a larger arena for males (50 × 50 × 50 cm) and in mating cages for females (Tecniplast 1284L model). Subsequently, 177 breeding pairs were established (generation 36: 37 P-line and 36 C-line pairs; generation 38: 52 P-line and 52 C-line pairs). One week after mating, males selected for the present study underwent a third hunting test (HT3). All hunting tests lasted 10 min and were preceded by a 3 min habituation period in the test arena before the cricket was introduced. Two days after HT3, the first intermale aggression test (IT1) was conducted, followed by the second test (IT2) two days later. Each intermale aggression test lasted 20 min (these three tests are described in detail in Materials and Methods section of the paper). The final analysed sample, which completed HT3, IT1, and IT2, comprised 45 C-line and 44 P-line males, corresponding to 10–12 males per replicate line and originating from 12–14 families per line, with 1–2 males sampled per family. After completion of behavioural testing, focal males were sacrificed for further analyses as part of the broader project.

### Supplementary tables

**Table S1**: Keywords and search logic used for a systematic search for studies examining whether selection for predatory behaviour influences intermale aggression, or whether selection for intermale aggression affects predatory behaviour. Searches were conducted in both Scopus and Web of Science databases. The search strategy combined three conceptual categories: (1) intermale or conspecific aggression, (2) predatory behaviour or hunting, and (3) evolution or artificial selection. Search output: The search returned 135 articles in Scopus and 183 articles in Web of Science. After screening, none directly tested whether selection for enhanced predatory behaviour leads to changes in intermale aggression.

| Concept | Keywords / Search terms |
|---|---|
| Intermale / Conspecific aggression | "intermale aggression" OR "conspecific aggression" OR "male–male aggression" OR "social aggression" OR "agonistic behaviour" OR "agonistic behavior" OR "intraspecific aggression" |
| Predatory behaviour / Hunting | predat* OR "predatory aggression" OR "predatory behavior" OR "predatory behaviour" OR hunt* OR "muricidal behavior" OR "muricidal behaviour" |
| Evolution / Selection | evolut* OR adapt* OR selection OR "experimental evolution" OR "artificial selection" |

**Table S2**: Definitions of behaviours and composite indices used in the cricket hunting and intermale aggression tests.

| Behaviour | Type | Definition |
|---|---|---|
| **Both cricket hunting intermale aggression tests** | | |
| Approach | Point | Directed movement towards the stimulus (cricket or opponent male) that reduced the distance between the focal vole and the stimulus, irrespective of whether it led to direct interaction or attack. |
| Resting | State | Periods during which the vole remained immobile and was not engaged in any other observable behaviour. |
| Observing | State | Stationary behaviour in which the vole visually attended to the stimulus without approaching or initiating interaction. |
| Exploration | State | General locomotor and exploratory activity not classified under any other behavioural category. It was not directly annotated during video scoring, but calculated as the residual active time, defined as the total test duration (600 s for the predatory test and 1200 s for the intermale test) minus the summed duration of all annotated state behaviours. |
| **Cricket hunting test** | | |
| Spotting | Point | Initial detection of the cricket, operationalised as the first clear orientation or visual attention towards the cricket. |
| Hunting | State | Active prey-directed behaviour including pursuit, rapid orientation, chasing, or lunging towards the cricket. |
| Capture | Point | Successful grasping of the cricket followed by killing it. |
| Eating | State | Consumption of the cricket following successful capture. |
| **Intermale aggression test** | | |
| Chasing | State | Rapid directed pursuit of one vole by another, resulting in a reduction in distance. |
| Boxing | State | Upright interaction involving forelimb striking or pushing between voles. |
| Wrestling | State | Close-contact physical interaction involving grappling or rolling. |
| Escaping | State | Movement away from another vole resulting in increased distance and avoidance of interaction. |
| Nosing or olfactory investigation | State | Sniffing or close-contact investigation using the nose directed towards the environment or another vole. |
| Any Aggression | State | Composite of chasing, boxing, and wrestling behaviours. |
| Supine posture | State | Posture in which the vole lies on its back for at least 3 seconds, typically associated with submissive or defensive responses. |
| Squatting posture | State | Low crouched posture with reduced movement, typically reflecting a non-engaging or defensive state. |
| Aggression index (duration) | - | Calculated as the sum of z-score standardised focal–opponent differences in $\log_{10}$-transformed behavioural durations, with aggressive components (boxing, wrestling, chasing) contributing positively and defensive components (escaping, supine posture, squatting) negatively. |
| Proactivity index | - | Calculated as the sum of focal–opponent differences in $\log_{10}$-transformed behavioural measures, with exploration and olfactory investigation contributing positively and resting duration and approach latency negatively. |
